# Augmented Super-Resolution Radial Fluctuations (aSRRF) Pushing the Limits of Structured Illumination Microscopy

**DOI:** 10.1101/2023.07.05.547885

**Authors:** Heng Zhang, Jianhang Wang, Luhong Jin, Yunqi Zhu, Yuting Guo, Meng Zhang, Yuhui Zhang, Zhixiong Wang, Yisun Su, Yicong Wu, Baohua Ji, Derek Toomre, Xu Liu, Yingke Xu

**Author notes:** Correspondence should be addressed to **Prof. Yingke Xu:** Department of Biomedical Engineering, 38 Zheda Road, Yuquan Campus, Zhejiang University, Hangzhou 310027, China. These authors contributed equally: Heng Zhang, Jianhang Wang and Luhong Jin.

## Abstract

Structured illumination microscopy (SIM) is a versatile super-resolution technique known for its compatibility with a wide range of probes and fast implementation. While 3D SIM is capable of achieving a spatial resolution of ∼120 nm laterally and ∼300 nm axially, attempting to further enhance the resolution through methods such as nonlinear SIM or 4-beam SIM introduces complexities in optical configurations, increased phototoxicity, and reduced temporal resolution.

Here, we have developed a novel method that combines SIM with augmented super-resolution radial fluctuations (aSRRF) utilizing a single image through image augmentation. By applying aSRRF reconstruction to SIM images, we can enhance the SIM resolution to ∼50 nm isotopically, without requiring any modifications to the optical system or sample acquisition process. Additionaly, we have incorporated the aSRRF approach into an ImageJ plugin and demonstrated its versatility across various fluorescence microscopy images, showcasing a remarkable two-fold resolution increase.

## Main

Super-resolution imaging (SRI) techniques, often referred to as ‘nanoscopes’, surpass the diffraction limit of light by employing either single molecule localization or patterned illumination, enabling the visualization of molecules and organelles in significantly higher detail within living cells^1,2^. Structured illumination microscopy (SIM) is among the most rapidly evolving SRI technique that uses the same multicolor probes as widefield or confocal microscopy, while enabling an approximately 2-fold increase in resolution compared to its widefield counterpart^3,4^. In contrast to single molecule switching that requires thousands of raw images to generate a super-resolved image, SIM achieves super-resolution (∼120 nm laterally and ∼300 nm axially) by typically using 15 sinusoidal patterned images in conjunction with specific reconstruction algorithms^3^. Therefore, SIM offers the advantage of reduced photobleaching, making it particularly suitable for live cell SRI. Advancements have been made in enhancing SIM resolution with nonlinear^5^ or 4-beam SIM^6^, developing algorithms to address SIM image artifacts^7–9^, and using deep learning techniques to minimize the number of raw images required for reconstruction^10,11^ or to achieve isotropic resolution^12^. Nevertheless, the strong desire to improve the spatial resolution of SIM persists, both with existing SIM platforms and even after implementing the aforementioned advancements.

Super-Resolution Radial Fluctuations (SRRF)^13^ is an innovative computational method that revolutionizes super-resolution reconstruction. By analyzing the radial and temporal fluorescence intensity fluctuations in a time-lapse image sequence, SRRF directly generates a super-resolved image. Unlike traditional methods that heavily rely on fluorophore detection and localization, SRRF eliminates the need for these processes, making super-resolution imaging more accessible and efficient^14^. Typically, SRRF requires the acquisition of 100-1000 raw time-lapse images to reconstruct a single high-resolution image. With the integration of GPU based platform, SRRF-Stream allows for real-time super-resolution reconstruction and visualization of dynamics in live cells^15^. It capitalizes on subtle frame-to-frame intensity variations, with theoretical potential to improve the resolution of various input data. Recently, an enhanced version called eSRRF (enhanced-SRRF), evolved from SRRF, can automatically select the minimal number of frames for optimal SRRF reconstruction, and is compatible with total internal reflection fluorescence (TIRF), spinning-disk confocal (SDC), and light-sheet microscopy^16^.

However, the requirement of a large quantity of raw images (e.g. ∼5000 frames for a 3D stack) for classic SRRF or eSRRF reconstruction poses limitations in rapid 3D imaging and leads to issues of photobleaching and photodamage. To overcome these challenges, here we aimed to significantly enhance the speed of SRRF and apply it to SIM for faster and higher-resolution imaging of live cells. For this purpose, we proposed a novel method called aSRRF (augmented SRRF), as illustrated in Figs. 1a-d. Importantly, aSRRF requires only a single input image and employes *in-silico* simulations along with Gaussian distribution properties of individual pixels to generate a synthetic time-lapse image sequence, known as image augmentation, for subsequent SRRF super-resolution reconstruction (see Methods). This approach greatly reduces light exposure and increases temporal resolution for live cell super-resolution imaging. Our results provide compelling evidence that the implementation of aSRRF effectively enhances both the lateral and axial resolutions of SIM by ∼2-fold. This substantial improvement ultimately leads to a remarkable resolution of ∼50 nm, particularly when applied to state-of-the-art 3D SIM with an initial resolution of approximately 120 nm isotopically^6^.

**Figure 1.**
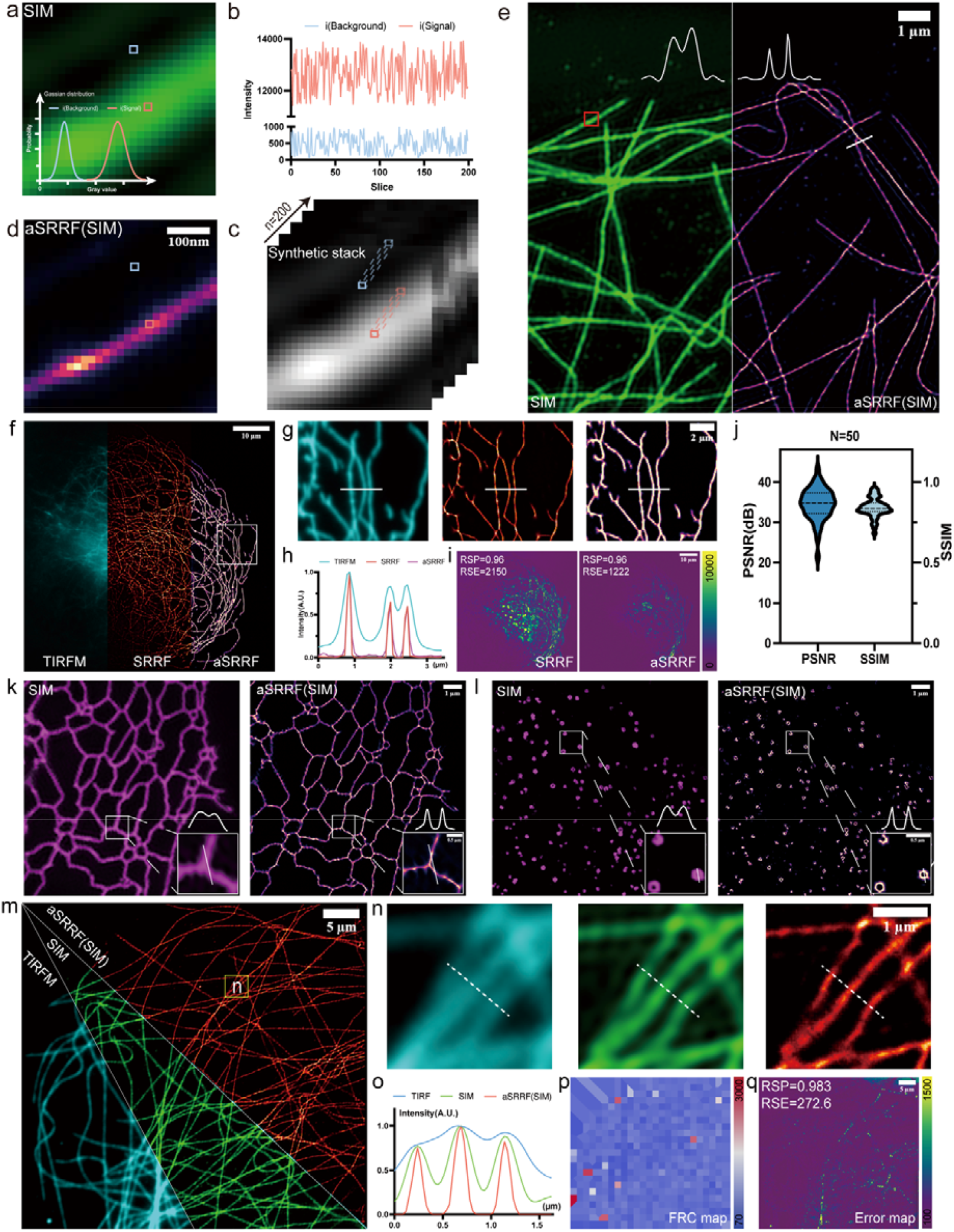
Schematic of aSRRF method and its performance in enhancing resolution in TIRFM and SIM. (a-d) Schematic illustration of aSRRF. (a) SIM image of fluorescently labeled microtubules. Displayed are higher-magnification view of regions indicated by the red box in (e). (b) Fluctuation map of individual pixels (red: fluorescence signal; blue: noise) sampled from distinct Gaussian distributions. (c) Augmented image stack consisting of 200 frames generated from (a). (d) aSRRF reconstruction of (c). (e) Comparative view of fluorescently labeled microtubules between SIM (left) and aSRRF reconstruction (right). The white curves overlaid on the images represent the fluorescence intensity distribution at the locations indicated by the white solid line. (f) Image of microtubules visualized with TIRFM (left), subsequently processed with classic SRRF (middle) or aSRRF (right). (g) Enlarged regions from the white box in (f). From left to right are the results of TIRFM, classic SRRF, and aSRRF, respectively. (h) Normalized intensity profiles of microtubules along the white lines indicated in (g). (i) Error maps comparing SRRF and aSRRF reconstructions to the original TIRFM image. (j) Quantitative analysis with PSNR (34.56͹±͹3.9, n = 50 cells) and SSIM (0.84͹±͹0.06, n = 50 cells) for aSRRF reconstruction using classic SRRF reconstruction as ground truth. (k-l) Images of ER (k) and CCPs (l), comparing SIM and aSRRF reconstruction. Inserts are the magnified views of white regions. The white curves displayed above the inserts represent line intensity profiles along the white lines, indicating the increased resolution achieved with aSRRF. (m) Microtubule structures visualized in TIRFM, SIM, and aSRRF reconstructions from SIM. (n) Enlarged regions from the white box in (m), comparing TIRFM (left), SIM (middle), and aSRRF reconstruction (right). (o) Normalized intensity profiles along the white dashed lines in (n). (p) FRC map that represents the resolution of aSRRF reconstruction (blocks=25). (q) Error map illustrates the reconstruction error of aSRRF compared to SIM reconstruction. Scale bars: (a)(c)(d), 100 nm; (e)(k)(l)(n), 1 μm; (f), 10 μm; (g), 2 μm; (m), 5 μm.

The schematic diagram of aSRRF is illustrated in Figs. 1a-d. For each pixel in the given image, a unique Gaussian distribution can be determined (Fig. 1a & Methods). Next, 200 values are randomly chosen from the Gaussian distribution to generate fluctuations on individual pixels (Fig. 1b). By repeatedly processing each pixel within the image, an augmented synthetic stack is generated (Fig. 1c). Finally, the synthetic image stack was utilized as input for the subsequent classic SRRF reconstruction (Fig. 1d). Through our investigation of fluorescence fluctuations, we have found that the Gaussian distribution provides slightly better performance in aSRRF reconstruction compared to the Uniform distribution, when evaluated using metrics such as peak signal-to-noise ratio (PSNR) and structural similarity (SSIM) (Fig. S1). Furthermore, we have demonstrated that aSRRF has robust performance in images with low signal-to-noise ratio (SNR), producing comparable results to classic SRRF reconstruction (Fig. S2).

To provide compelling experimental evidence that aSRRF is effective in improving resolution, we conducted initial validation using simulated diffraction-limited spots. The outcomes showed that aSRRF successfully resolved spots with a separate of 60 nm, which would otherwise be indistinguishable using SIM (Fig. S3). Then, we imaged COS-7 cells (n=20) with fluorescently labelled endoplasmic reticulum (ER), and compared the results obtained from three different approaches: (1) diffraction-limited image, which was generated by averaging 9 raw SIM images; (2) aSRRF-processed diffraction-limited image, where the aSRRF algorithm was applied to the average of the raw SIM images; (3) SIM image, which was obtained through conventional SIM reconstruction. Comparison results suggest that the aSRRF reconstruction enhances resolution (77.4 ± 8 nm, *n*=20) relative to diffraction-limited images (272.6 ± 5 nm, *n*=20), which is comparable or even superior to SIM image (146.8 ± 8 nm, *n* =20), as evaluated by resolution-scaled Pearson’s coefficient (RSP) and resolution-scaled error (RSE) measurements (Fig. S4).

We next applied the aSRRF algorithm to enhance SIM imaging, with the goal of doubling the SIM resolution to below 60 nm. We conducted SIM imaging of nanorulers (∼60-80 nm) with different geometries. The results clearly demonstrated that aSRRF was able to resolve these structures that would otherwise be indistinguishable using SIM alone (Fig. S5). We further evaluated the performance of aSRRF in enhancing the resolution of TIRFM and SIM imaging using biological samples. Visually, the application of aSRRF clearly distinguished microtubules that appeared fused or indistinguishable in SIM images, providing strong support for its effectiveness in resolution enhancement (Fig. 1e).

To validate the reliability of aSRRF reconstruction, we conducted a comparative analysis with classic SRRF, which uses experimentally acquired time-lapse images as input. Intuitively, both SRRF and aSRRF consistently improved TIRFM resolution (Figs. 1f-h). The quantitative restoration error map^17^ highlights the regions of discrepancy between the SRRF/aSRRF and original TIRFM image, confirming that aSRRF achieves comparable results to classic SRRF (Fig. 1i). Importantly, aSRRF only requires a single image, whereas SRRF requires about 100∼200 images, which could introduce temporal blurring due to potential sample motion or changes during the acquisition process. In addition, a statistical analysis comparing the aSRRF reconstruction to the reference classic SRRF indicated that aSRRF produced reliable results as assessed by PSNR and SSIM (Fig. 1j).

Similarly, we found that aSRRF also enhances the resolution of SIM in imaging of both ER (Fig. 1k) and doughnut-shaped clathrin-coated pits (CCPs)^18^ (Fig. 1l) structures. The side-by-side comparison results (Figs. 1m, n) demonstrated that aSRRF achieves the best performance in high-resolution imaging of microtubules (Fig. 1o). The Fourier Ring Correlation (FRC)^17^ map of aSRRF clearly illustrated its effectiveness in resolution enhancement (Fig. 1p). The restoration error map for the aSRRF versus the original SIM image, along with the global quality metrics RSP and RSE measurements, showed the high reliability of aSRRF reconstruction (Fig. 1q). In contrast to deep learning-based methods for 2-3 fold resolution enhancement^19^, aSRRF offers the advantage of being structure-independent and does not require any training data, making it a user-friendly and easily implementable solution.

To demonstrate the capabilities of aSRRF in live cell multi-color 3D super-resolution imaging, we conducted a thorough analysis comparing its resolution enhancement with regular 2D and 3D SIM imaging techniques. Our results revealed that aSRRF effectively captured organelle interactions, allowing for clear visualization of the elongation of ER tubules along the microtubules (marked white arrows in Fig. 2a). Furthermore, we showcased the application of aSRRF in the state-of-the-art 3D SIM imaging with an isotropic resolution of ∼120 nm^6^, demonstrating its versatility and effectiveness in super-resolution reconstruction (Fig. 2b). Specifically, the application of aSRRF results in an approximate 2-fold increase in lateral resolution from 108 nm to 55 nm (Figs. 2b, d), and axial resolution as well, from 107 nm to 53 nm (Figs. 2c, e, f). Addtinally, aSRRF enables the resolution of the separation process of lysosomes that would otherwise be indistinguishable by 3D SIM imaging, even after employing deep learning technique (Fig. S6). Collectively, these results demonstrated that aSRRF is capable of pushing the limits of SIM imaging to ∼50 nm, enabling new insights into the study of biological dynamics at sub-100 nm scale.

**Figure 2.**
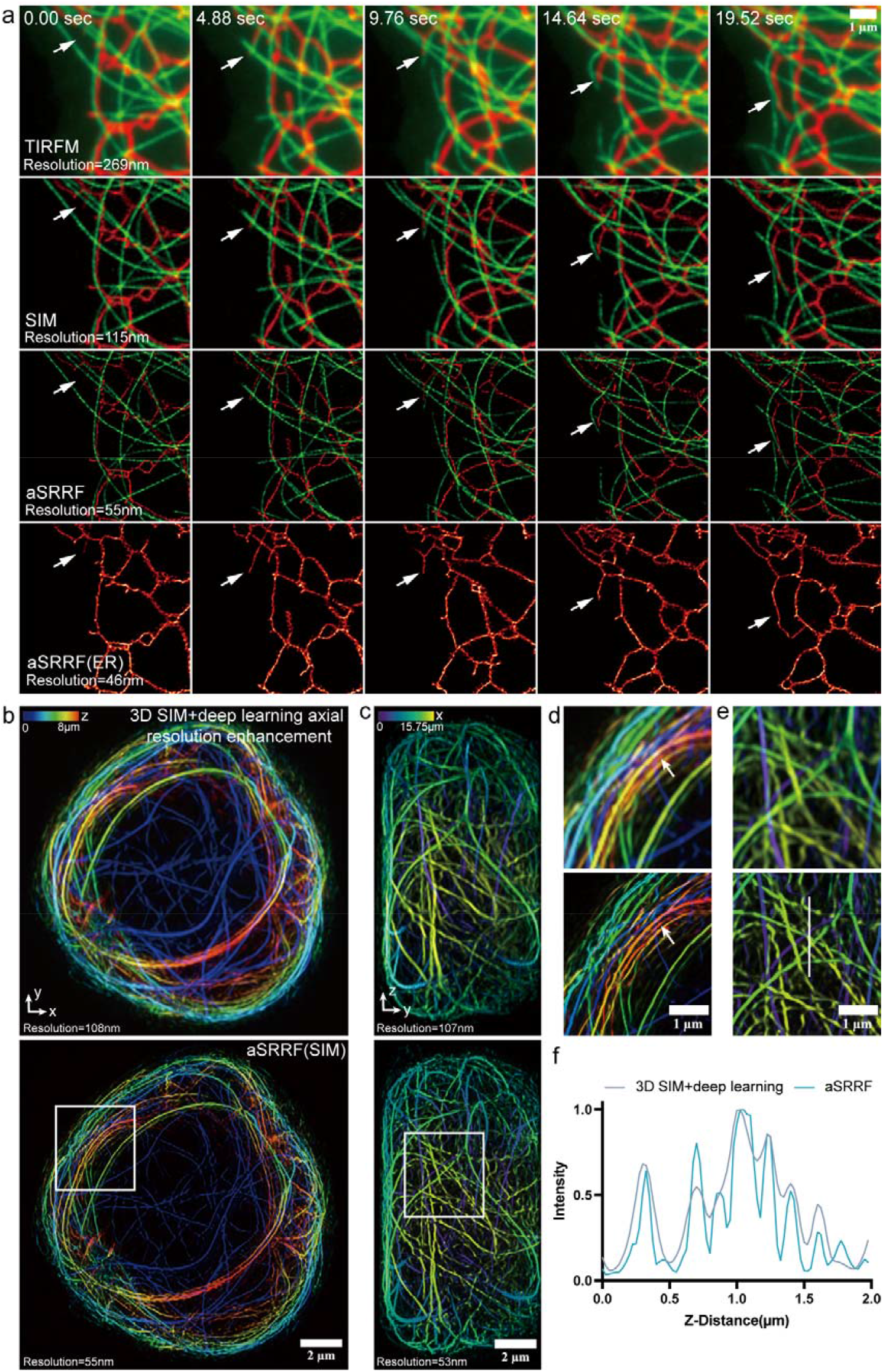
The aSRRF doubles both lateral and axial resolutions in multi-color live cell SIM imaging. (a) aSRRF elevates the resolution of dual-color live cell SIM imaging. The COS-7 cells were transfected with 3XmEmerald-ensconsin and mCherry-KDEL plasmids to label the microtubules and ER, respectively. Traction interactions between microtubules and ER can be visualized in aSRRF (marked by white arrowheads). (b, c) Lateral (b) and axial (c) view of a Jurkat T cell expressing microtubules, comparing 3D SIM with enhanced axial resolution through deep learning (top) and subsequently processed by aSRRF (bottom). Images are depth coded as indicated. (d, e) Enlarged regions of the white boxes indicated in (b) and (c), respectively. The white arrows in (d) highlight the resolution enhancement of aSRRF in differentiating the microtubule network. (f) Normalized intensity profile of the white solid line in (e). Scale bars: (a)(d)(e), 1 μm; (b)(c), 2 μm.

Our proposed aSRRF method offers a practical and cost-effective approach for super resolution imaging. It utilizes only a single image to simulate fluorescence fluctuations, employs image augmentation and reconstruction techniques, and eventually enhances both spatial and temporal resolution for live cell super-resolution imaging. This method is also compatible with other imaging modalities, such as epifluorescence microscopy, TIRFM (Fig. 1), multiview confocal super-resolutution microscopy^20^ (Fig. S7), and light-sheet microscopy^21^ (Fig. S8). We anticipate that aSRRF will have widespread applications in live cell super-resolution fluorescence imaging.

## Data availability

The MATLAB codes of aSRRF and the developed ImageJ plugin for aSRRF reconstruction are available for download at https://github.com/XuLabZJU/ImageStackAugmentation. Some source images were uploaded at Github (Fig. 1m, Fig. S3a, Fig. S5f) for user validation. Other relevant data are available from the corresponding author upon request, due to the size limits.

## Acknowledgments

This work was supported by the National Key Research and Development Program of China (2021YFF0700305), Zhejiang Provincial Natural Science Foundation (LZ23H180002), the National Natural Science Foundation of China (62105288 and 22104129), Zhejiang University K.P.Chao’s High Technology Development Foundation (2022RC009), the Fundamental Research Funds for the Central Universities (226-2023-00091) and the Zhejiang University Education Foundation Global Partnership Fund. Z.W., Y.S. and Y.W. acknowledge support from the intramural research program of the National Institute of Biomedical Imaging and Bioengineering, NIH.

## Author contributions

Y.X., H.Z. and L.J. conceived and initiated the project. H.Z., J.W., L.J. and Y.Z. performed experiments. Y.G. performed live cell SIM imaging. M.Z. and Y.Z. designed and performed the light sheet imaging. Z.W., Y.S., and Y.W. conducted the 3D SIM and multi-view confocal SIM imaging and analysis. Y.W., B.J., D.T. and X.L. provided key inputs. Y.X. supervised the project.

## Competing interests

The authors declare no competing financial interests.

## Materials and Methods

### Cell culture

COS-7 cells were cultured in high-glucose Dulbecco’s modified Eagle’s medium (DMEM; Thermo Fisher Scientific) supplemented with 10% (v/v) fetal bovine serum (FBS; Corning), 1% (v/v) penicillin/streptomycin (Thermo Fisher Scientific) and 2⍰mM L-alanyl-L-glutamine (GlutaMAX™, Thermo Fisher Scientific) at 37⍰°C in a 5% CO_2_ incubator.

### Sample preparations

COS-7 cells cultured on 35 mm glass bottom MatTek dishes were fixed with 3% glutaraldehyde and 0.25% Triton-X for 15 min. After fixation, the cells were immunostained with a primary antibody against α-tubulin and a secondary antibody conjugated with Alexa Fluor 488 as previously described^22^. For live cell imaging of ER and microtubules, the COS-7 cells were transfected with plasmids encoding tubulin (3×mEmerald-ensconsin) and ER (mCherry-KDEL) by using Lipofectamine 3000 (Invitrogen) according to the manufacturer’s protocol^17^. The cells were imaged 24 hours post-transfection in KRBH buffer (pH 7.4), within a microscope stage top micro-incubator (OKO Lab) maintained at 37°C and 5% CO_2_. Commercial fluorescent samples (theGATTA-SIM 80 and the Argo-SIM V2.0 slide, Argolight) with 80 nm fluorescing beams distances and ground truth patterns consisting of fluorescing double line pairs (spacing from 0 nm to 390 nm) were used for comparison of resolution between different imaging modalities.

The mice esophagus tissue sections were prepared following the protocol previously published^20^. Specifically, they were obtained from 8-week old male C57BL mice provided by the section on Molecular Morphogenesis at the Eunice Kennedy Shriver National Institute of Child Health and Human Development (NICHD). The animal housing facilities maintained a 12-hour light cycle, a temperature range of 70-74 °F, and a humidity range of 30%-70%. The animal protocols were approved by the Animal Care and Use Committees in NICHD. To prepare the tissue sections, the esophagus tissue from C57BL mice were fixed in 4% PFA/PBS at 4°C overnight. The fixed tissue was then subjected to a series of sucrose/water solution (Sigma, S0389) washes with gradually increasing concentrations (10%, 20%, and 30%) for 3 hours each at room temperature (RT). Following this, the tissue was equilibrated in 30% sucrose solution at 4°C overnight. The tissue was subsequently cut into cubes using a razor blade and immersed in Optimal Cutting Temperature solution (O.C.T, Fischer Scientific NC9806257) within a mold (Fischer Scientific, 22-363-552). The tissue cubes were then frozen in a solution of pure ethanol and dry ice for 10 min. Frozen tissue was sectioned to 25 μm thicknesses using a Leica CM1850 cryostat and stored in 1X PBS. For immunostaining, the esophagus tissue sections underwent antigen retrieval by incubating in an antigen retrieval buffer (Abcam, ab93684) at 60°C for 1 hour. Following this step, the tissue sections were incubated overnight at 4°C with a primary antibody solution containing 1 μg/mL of mouse-α-Tubulin in 0.1% Triton-X/PBS. Afterward, the sections were washed in 0.1% Triton-X/PBS for 1 hour at RT and then incubated with a secondary antibody solution comprising 1 μg/mL of α-mouse-JF549 and 10 μM of phalloidin-Alexa Fluor 488 (Thermo Fisher Scientific, A12379) in 0.1% Triton-X/PBS for 1 hour at room teperature. The stained tissue sections were washed in 0.1% Triton-X/PBS at RT for 1 hour and finally mounted on poly-L-lysine coated cover glasses (VWR, 48393-241) for triple-view line confocal super-resolution imaging^20^.

### Microscopes

TIRFM imaging was performed on a customized ring-illumination TIRFM system^22^ equipped with 488 nm (50 mW, Coherent) and 561 nm (100 mW, Coherent) laser lines and a 100× NA 1.49 TIRF objective (Olympus UAPON100XOTIRF). Fluorescence emission was collected onto an EMCCD camera (Andor iXon Life 888), yielding a pixel size of 133 nm. A custom-developed SIM system was used for dual-color 2D live cell imaging^18^, which equipped with 488 nm (500 mW, Coherent) and 561 nm (500 mW, Coherent) laser lines and a 100× NA1.7 TIRF objective (Olympus APON 100XHOTIRF). High speed sCOMS camera (Hamamatsu Orca Flash 4.0 v3) was used for dynamic imaging, and the nine raw images consisting of 3-orientaion × 3-phase for each time point were reconstructed into a super-resolution image based on the previous published algorithm^23^. A home-built SIM system equipped with 1.2 NA water immersion lens was used for conucting 3D SIM imaging of microtubules and LAMP1-labelled lysosomes^6^. The acquired 3D SIM results were subsequently processed using a previously published deep learning method^20^, leading to an isotropic resolution of ∼120 nm. For imaging the mouse esophageous tissue section, a multi-view line confocal structured illumination microscopy setup with 0.8, 0.8, and 1.2 NA water immersion lenses was employed, as described in a previous publication^20^. Light-sheet imaging was performed on a custom-built double-ring Bessel light-sheet microscope^21^.

### aSRRF reconstruction

The effectiveness of aSRRF is due to its ability to simulate the temporal fluctuations of fluorescent molecules at the single pixel level, which enables it to improve the resolution of images obtained from microscopes that have difficulties in capturing temporal fluctuations of fluorescent molecules in 100-200 image sequences. Classic SRRF algorithm relies on the temporal fluorescence intensity fluctuations to improve image resolution through subpixel refinement to distinguish between signal and noise. By simulating the temporal fluctuations of individual pixels, aSRRF can improve the resolution of imaging by only utilizing one single image.

The simulation process of Gaussian and Uniform distribution in a single pixel can be represented by the following equations:

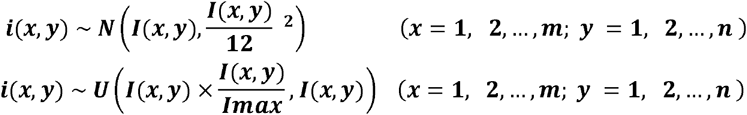

Here *i(x, y)* denotes the pixel value of x rows and y columns in the image. N denotes Gaussian distribution and *U* denotes Uniform distribution. *Imax* represents the maximum pixel value in the image. *i(x, y)* represents the pixel value at the current coordinate position, and *m, n* denote the width and height of the image, respectively. For image augmentation, 200 values are chosen randomly from the Gaussian or Uniform distribution to generate the fluorescence intensity fluctuations on single pixels. By repeatedly processing each pixel within the image, an augmented synthetic stack consists of 200 frames is obtained. Finally, the SRRF plugin in Fiji/ImageJ (NIH) is used to process the generated image stack, by using radiality magnification and temporal radiality average as temporal analysis functions. Different number of synthetic images were generated and the performance of aSRRF reconstruction has been quantitatively analyzed. As shown on Fig. S2, synthetic generation of 200 images are sufficient to achieve resolution enhancement even in low SNR conditions. We have also conducted extensive validations on time-lapse image sequences and have found that most of the pixels, regardness of signal or noise, display distinct Gaussian distributions (Supplementary Video S1). Therefore, we modeled the fluctuations of each pixel by using Gaussian distribution, in which the mean is the current pixel value and the variance is the current pixel value divided by 12 based on experimental optimizations. A comparison of the Gaussian and Uniform distributions is also shown in Supplementary Video S2.

### Quantification the performance of aSRRF

The parameters PSNR, SSIM, RSP and RSE are used to evaluate the performance of aSRRF reconstruction. The image resolution was measured by ImageDecorrleationAnalysis plugin in Fiji/ImageJ (NIH) with default settings^24^. PSNR and SSIM were calculated as the equations shown below:

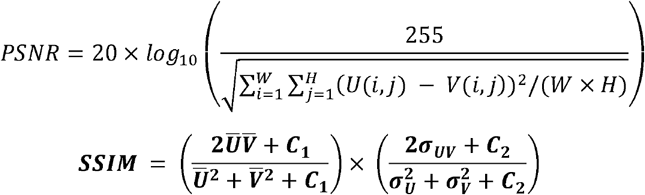

Here W and H represent the width and height of the reference image (e.g. Classic SRRF reconstruction results). U and V represent the SRRF and aSRRF reconstruction, respectively. Ū and 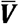 denote the averages of U and V, whereas **σ**_**U**_ and **σ** _**V**_ denote the variances of U and V. **σ** _**UV**_ is the covariance of U and V. The parameters C1 and C2 are positive constants used to stabilize each term in the equation ***C***_***1***_ ***= (k***_***1***_***L)*** ^***2***^, ***C***_***2***_ ***= (k***_***2***_***L)*** ^***2***^. L represents the dynamic range of the pixel values, and k1 and k2 have default values of 0.01 and 0.03 respectively. The code for image quantification was written in Python.

The super-resolution quantitative image rating and reporting of error locations (SQUIRREL) method was applied to evaluate the global image quality. SQUIRREL calculates the global RSE and RSP values at the super-resolution image against the diffraction-limited image. Moreover, the Fourier ring correlation map, calculated by the SQUIRREL plugin in Fiji/ImageJ (NIH), illustrates the resolution distribution in each block.

## Notes

https://github.com/XuLabZJU/ImageStackAugmentation

### Competing Interest Statement

The authors have declared no competing interest.

### Summary of Updates

Figure Format Change

## References

1. Sahl, S. J., Hell, S. W. & Jakobs, S. Fluorescence nanoscopy in cell biology. Nat. Rev. Mol. Cell Biol. 18, 685–701 (2017).

2. Schermelleh, L. et al. Super-resolution microscopy demystified. Nat. Cell Biol. 21, 72–84 (2019).

3. Gustafsson, M. G. L. Surpassing the lateral resolution limit by a factor of two using structured illumination microscopy. J. Microsc. 198, 82–87 (2000).

4. Wu, Y. & Shroff, H. Faster, sharper, and deeper: structured illumination microscopy for biological imaging. Nat. Methods 15, 1011–1019 (2018).

5. Rego, E. H. et al. Nonlinear structured-illumination microscopy with a photoswitchable protein reveals cellular structures at 50-nm resolution. Proc. Natl. Acad. Sci. 109, E135–E143 (2012).

6. Li, X. et al. Three-dimensional structured illumination microscopy with enhanced axial resolution. Nat. Biotechnol. 1–13 (2023) doi:10.1038/s41587-022-01651-1.

7. Wen, G. et al. High-fidelity structured illumination microscopy by point-spread-function engineering. Light Sci. Appl. 10, 70 (2021).

8. Smith, C. S. et al. Structured illumination microscopy with noise-controlled image reconstructions. Nat. Methods 18, 821–828 (2021).

9. Huang, X. et al. Fast, long-term, super-resolution imaging with Hessian structured illumination microscopy. Nat. Biotechnol. 36, 451–459 (2018).

10. Jin, L. et al. Deep learning enables structured illumination microscopy with low light levels and enhanced speed. Nat. Commun. 11, 1934 (2020).

11. Xypakis, E. et al. Deep learning for blind structured illumination microscopy. Sci. Rep. 12, 8623 (2022).

12. Guo, M. et al. Rapid image deconvolution and multiview fusion for optical microscopy. Nat. Biotechnol. 38, 1337–1346 (2020).

13. Gustafsson, N. et al. Fast live-cell conventional fluorophore nanoscopy with ImageJ through super-resolution radial fluctuations. Nat. Commun. 7, 12471 (2016).

14. Culley, S., Tosheva, K. L., Matos Pereira, P. & Henriques, R. SRRF: Universal live-cell super-resolution microscopy. Int. J. Biochem. Cell Biol. 101, 74–79 (2018).

15. Browne, M. et al./person-group>. Real time multi-modal super-resolution microscopy through Super-Resolution Radial Fluctuations (SRRF-Stream). in Single Molecule Spectroscopy and Superresolution Imaging XII (eds. Gregor, I., Gryczynski, Z. K. & Koberling, F.) 42 (SPIE, 2019). doi:10.1117/12.2510761.

16. Laine, R. F. et al. High-fidelity 3D live-cell nanoscopy through data-driven enhanced super-resolution radial fluctuation. bioRxiv 2022.04.07.487490 (2022) doi:10.1101/2022.04.07.487490.

17. Culley, S. et al. Quantitative mapping and minimization of super-resolution optical imaging artifacts. Nat. Methods 15, 263–266 (2018).

18. Qiao, C. et al. Evaluation and development of deep neural networks for image super-resolution in optical microscopy. Nat. Methods 18, 194–202 (2021).

19. Qiao, C. et al. Rationalized deep learning super-resolution microscopy for sustained live imaging of rapid subcellular processes. Nat. Biotechnol. 41, 367–377 (2023).

20. Wu, Y. et al. Multiview confocal super-resolution microscopy. Nature 600, 279–284 (2021).

21. Zhao, Y. et al. Isotropic super-resolution light-sheet microscopy of dynamic intracellular structures at subsecond timescales. Nat. Methods 19, 359–369 (2022).

22. Jin, L. et al. High-resolution 3D reconstruction of microtubule structures by quantitative multi-angle total internal reflection fluorescence microscopy. Opt. Commun. 395, 16–23 (2017).

23. Gustafsson, M. G. L. et al. Three-Dimensional Resolution Doubling in Wide-Field Fluorescence Microscopy by Structured Illumination. Biophys. J. 94, 4957–4970 (2008).

24. Descloux, A., Grußmayer, K. S. & Radenovic, A. Parameter-free image resolution estimation based on decorrelation analysis. Nat. Methods 16, 918–924 (2019).

